# Quantifying trade-offs in the choice of ribosomal barcoding markers for fungal amplicon sequencing: a case study on the grapevine trunk mycobiome

**DOI:** 10.1101/2022.07.01.498490

**Authors:** Vinciane Monod, Valérie Hofstetter, Vivian Zufferey, Olivier Viret, Katia Gindro, Daniel Croll

## Abstract

The evolution of sequencing technology and multiplexing has rapidly expanded our ability to characterize fungal diversity in the environment. However, obtaining an unbiased assessment of the fungal community using ribosomal markers remains challenging. Longer amplicons were shown to improve taxonomic resolution and resolve ambiguities by reducing the risk of spurious operational taxonomic units. We examined the implications of barcoding strategies by amplifying and sequencing two ribosomal DNA fragments. We analyzed the performance of the full internal transcribed spacer (ITS) and a longer fragment including also a part of the 28S replicated on 60 grapevine trunk core samples. Grapevine trunks harbor highly diverse fungal communities with implications for disease development. Using identical handling, amplification and sequencing procedures, we obtained higher sequencing depths for the shorter ITS amplicon. Despite the more limited access to polymorphism, the overall diversity in amplified sequence variants was higher for the shorter ITS amplicon. We detected no meaningful bias in the phylogenetic composition due to the amplicon choice across analyzed samples. Despite the increased resolution of the longer ITS-28S amplicon, the higher and more consistent yields of the shorter amplicons produced a clearer resolution of the fungal community of grapevine stem samples. Our study highlights that the choice of ribosomal amplicons should be carefully evaluated and adjusted according to specific goals.

## Introduction

Fungi occur in nearly all environments, are highly diverse and can form tight associations with other organisms as pathogens or mutualists (1). The mycobiome associated with plants has important implications for agricultural ecosystems (2). Vascular diseases affecting plant stems including xylem and phloem are often difficult to diagnose or the causal agent is not yet known (3–6). Surveys of fungal communities (*i*.*e*., the mycobiome) have become key tools to understand how environmental and temporal factors influence species compositions associated with diseases (7, 8). The evolution of sequencing technology and multiplexing has rapidly expanded our ability to characterize fungal diversity in many environments (9). However, the implementation of molecular tools to establish unbiased mycobiome surveys remains challenging (9). Early impediments of surveying fungal diversity included the need to culture species, which creates significant biases in the estimation of community compositions (10, 11). Next-generation sequencing (NGS) technology has enabled the sequencing of taxonomically informative loci (*i*.*e*., barcoding) to reproducibly determine community structures and species diversity (1). Second generation sequencing techniques *(i*.*e*., short read sequencing) can generate deep coverage amplicon datasets (12). Read length constraints limit amplicons to *ca*. 550 base pairs using an overlapping paired-end design (13).

The fungal nuclear ribosomal internal transcribed spacers (ITS) 1 and 2, in addition to the 5.8S subunit is the prevalent barcoding locus for fungi (14). With a typical amplicon length of 400-600 bp, the locus is compatible with second generation sequencing read length limitations (15). However, several studies have highlighted potential shortcomings of relying on short barcoding markers (16, 17). Targeting longer amplicons has been made possible by third generation sequencing technology. Longer amplicons were shown to improve taxonomic resolution (18) and resolve ambiguities by reducing the risk of spurious operational taxonomic units (OTU) (19–21). An important factor in establishing long read amplicon sequencing are error-correction approaches such as the PacBio circular consensus sequences (CCS). CCS drastically reduces base-calling errors through multiple sequencing passes on the same molecules (19). Importantly, CCS provides a per-base accuracy comparable to that of short-read sequencing (22, 23). However, the choice of the locus, challenges in amplifying longer fragments and downstream analyses remain important considerations.

Comparisons between long and short amplicon studies of fungal communities show that long amplicons have typically lower taxonomic coverage in sequence databases (24). However, longer reads can improve taxonomic resolution (25, 26). It raises the question whether targeting a longer amplicon (*i*.*e*., full ITS or full ITS + flanking rRNA subunit regions) can provide sufficient sequence read depth per sample to accurately capture relevant differences in species richness and community composition (27). Use of the full ITS region combines the benefits of capturing both ITS1 and ITS2 subregions (20). Targeting the full 5.8S rRNA gene provides improvements for fungal identification notably the precision of genus-level identification because of a much lower substitution rate compared to ITS1 or ITS2 (1).

The higher taxonomic resolution given by the full-ITS can be used for strain-level identifications (28, 29). Tedersoo et al. (26) have shown that the identification rate was 33% higher at genus rank when using full-length ITS sequences compared to using either ITS1 or ITS2. Longer sequences that combine the ITS with a portion of the small (18S) or large (28S) nuclear ribosomal subunits can also facilitate taxonomic assignments at the family or order level (17). In addition, more than 50% of taxonomically unassigned fungal ITS sequences, i.e., sequences corresponding to species belonging to underrepresented or not yet represented groups in sequence databases can be identified at least at the divisional level by adding flanking rDNA regions (26). Assessing the impact of using one or more ribosomal DNA regions for fungal identification should inform decisions about the design of fungal community and barcode studies.

As a model to assess fungal barcoding amplicon suitability, we focused on the complex grapevine trunk mycobiome. The grapevine trunk is inhabited by various fungal species from different taxonomic and functional groups (30). Determining fungal diversity is of high interest because various dieback diseases are thought to be caused by fungal pathogens constituting severe threats to vineyards worldwide with substantial economic consequences (31). Grapevine is subject to a complex set of interacting pathogenic or commensal microorganisms (2). Despite significant efforts over the past three decades, the causal agents among the grapevine trunk microbiome and the outbreak dynamics are poorly understood (30, 32). Expansive characterizations of the fungal diversity present in grapevine trunks may help to identify one or more fungal species strongly associated with disease outbreaks. To date, the grapevine mycobiome has been analysed largely based on culture-dependent approaches. Up to 159 OTUs were described using Sanger sequencing (31) and up to 259 OTUs with second generation NGS technology (30). Third generation long fragment sequencing has not been used to describe the grapevine mycobiome to our knowledge.

Here, we examine the implications of barcoding strategies in the context of the grapevine trunk mycobiome using third generation sequencing technology. We amplified and sequenced two fragments to analyse their performance in characterizing the grapevine trunk mycobiome: the full ITS (14) and a longer fragment composed of the full ITS and a part of the LSU to assess the impact on taxonomic resolution and phylogenetic coverage. We analysed whether the length of the target amplicon influences the sequencing depth and whether the sequencing depth correlates with the detected diversity. Finally, we examined if the amplicon choice influenced the detected fungal diversity.

## Results

### Amplicon sequencing and read recovery

A total of 60 grapevine plants located in a single plot were sampled at the grafting point using wood cores (Figure 1A). The prevalence of grapevine trunk diseases (GTD) among the 60 randomly selected plants was 14% in the sampling year. DNA extracted from the 60 wood samples could be successfully amplified in 59 and 50 samples with the primer pairs ITS1F-ITS4 (ITS) and ITS1F-LR5 (ITS-28S) respectively. One ITS and 10 ITS-28S amplicon samples were not sequenced because of a too low PCR product yield after amplification. A total of 382,672 PacBio CCS reads were successfully demultiplexed using the 8 bp barcode with 100% identity, generating 682-15’663 reads per sample for ITS (>4156 reads for 75% of the samples). For ITS-28S, 227’549 raw reads were demultiplexed, generating 707-22’288 reads per samples (>1876 reads for 75% of the samples). Sequencing depths were highly variable among samples for both loci (Figure 2A). A total of 90% of the reads were successfully demultiplexed for ITS and 89% for ITS-28S. In proportion, the ITS locus accounted for 63% of the total number of reads. The number of reads per sample between the two amplicons was not correlated (Figure 2A). This shows that the equimolar pooling of the PCR products was successful in balancing sequencing yields. Sequence lengths were comparable to the expected PCR amplicon sizes. The average sequence length was 629 pb for the ITS and 1526 bp for the ITS-28S (Figure 2B).

**Figure 1:**
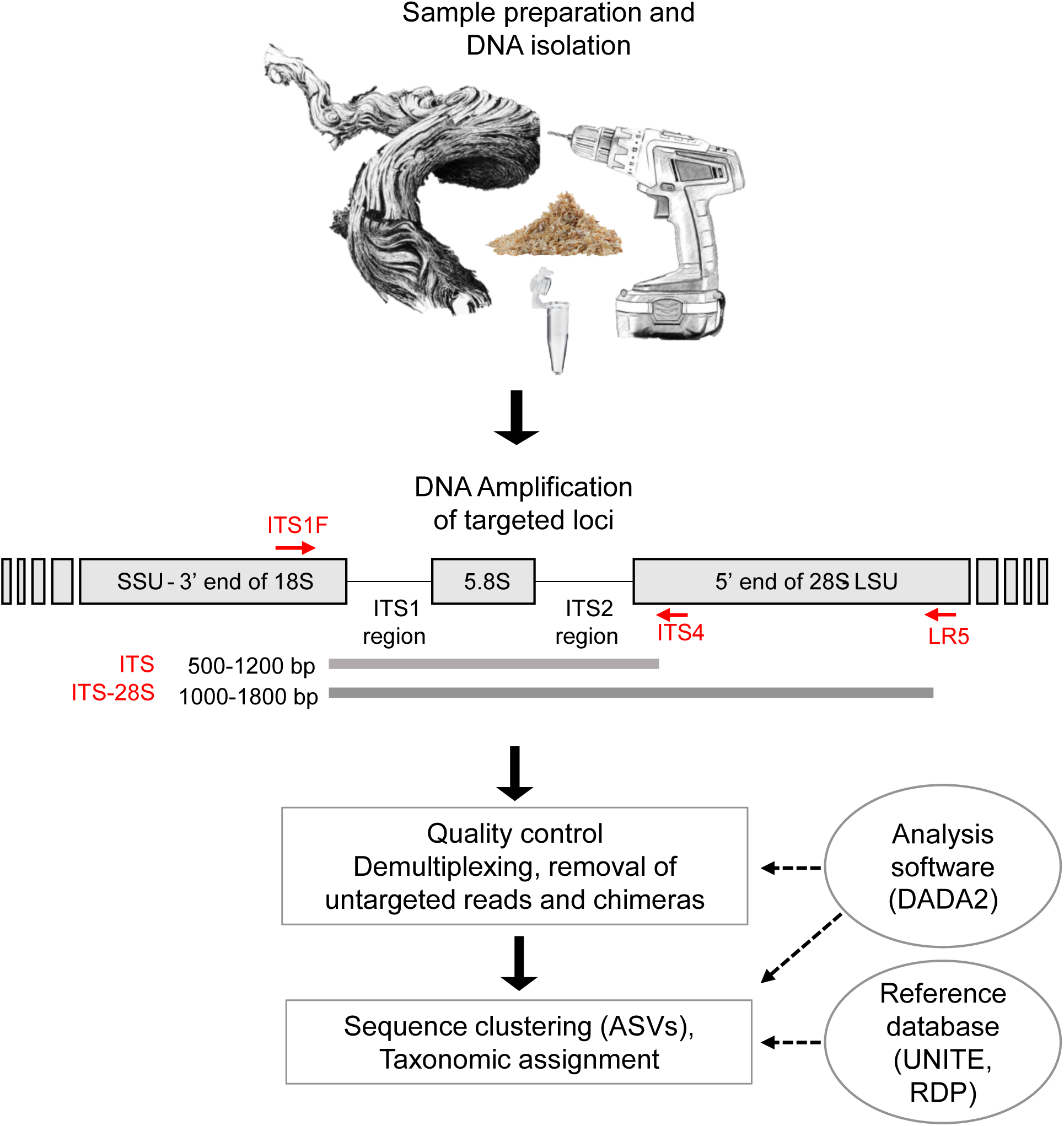
Trunk sample collection and barcoding loci. Sampling method used to extract wood cores from the grafting point of vine plants (*n* = 60) of a single vineyard before proceeding to DNA extraction. Genomic regions targeted for the amplification: internal transcribed spacer (ITS) and ITS - large subunit (LSU/28S) using the ITS1F − ITS4 and ITS1F − LR5 primer pairs, respectively. Bioinformatics workflow to filter and trim sequences following the DADA2 method. Inference of amplicon sequence variants (ASVs) from sequencing data and taxonomic assignments of each ASVs.

**Figure 2:**
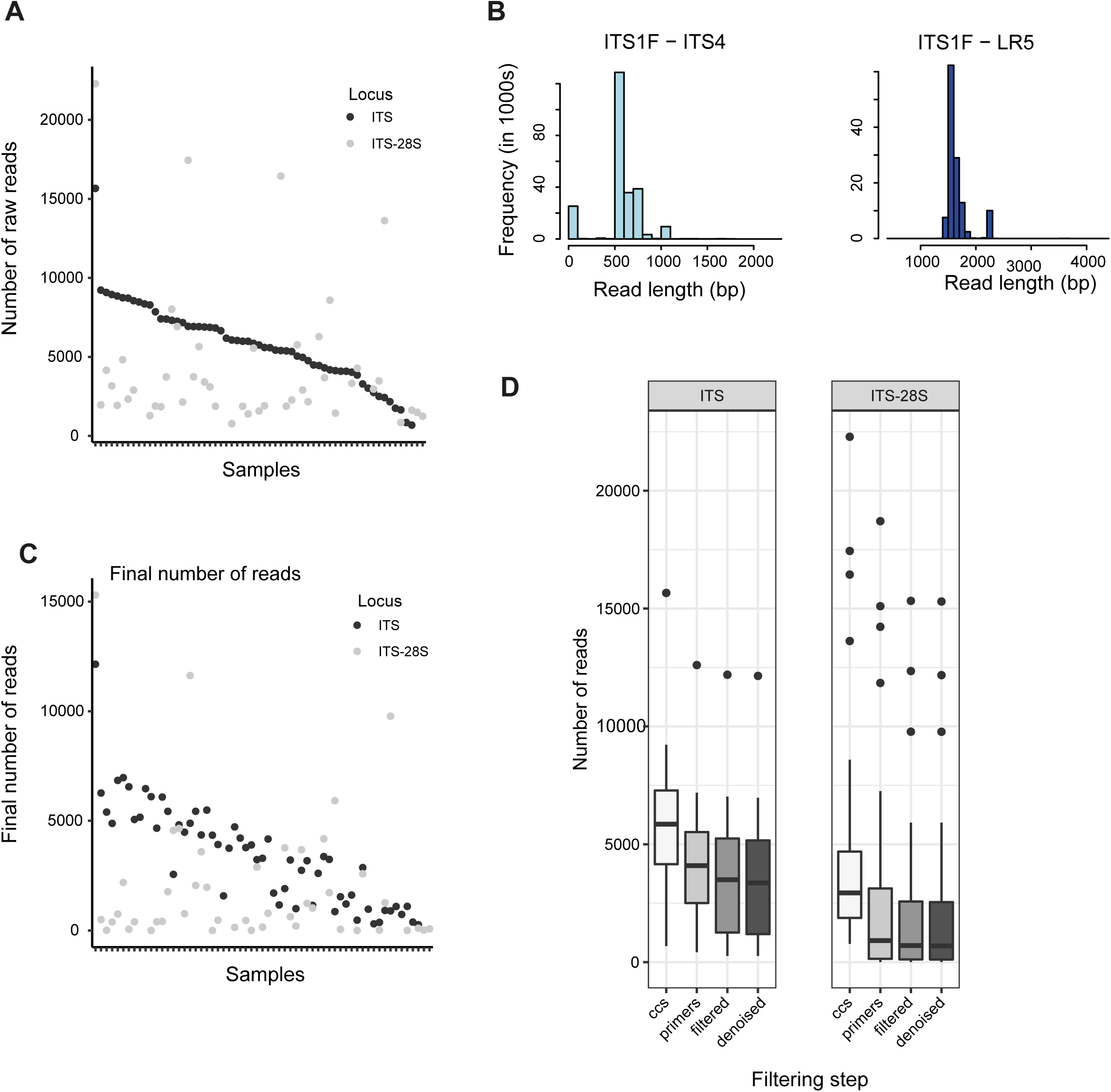
Circular consensus sequencing (CCS) analyses of two ribosomal amplicons. A) Number of raw CCS reads obtained for each of the 109 samples for the internal transcribed spacer (ITS) and ITS - large subunit (28S) amplicons amplified using the ITS1F − ITS4 and ITS1F − LR5 primer pairs, respectively. Samples are ranked by raw read counts of the ITS amplicon. B) Distribution of raw read lengths for each of the two amplicons for all samples combined. C) Number of final read numbers for the two amplicons after all filtering steps. Samples are ranked by raw read counts of the ITS amplicon. D) Impact of individual filtering steps (dereplication, primer detection, filtering for amplicon length, denoising according to error detection and presence of chimeras) on the read counts per sample for each of the two amplicons.

De-replication according to detected flanking primer sequences is a critical step to ensure high-quality sequences but this step can also discard a substantial number of sequences. For ITS, 29% of reads were discarded at this step and 38% were discarded for the ITS-28S. Three samples from the ITS-28S saw no reads passing the de-replication as no matching primer sequences were detected. Read filtering for amplicon length showed disparities between the ITS and ITS-28S datasets with 70% of reads retained for ITS and 61% retained for the ITS-28S (Figure 2D). Error detection and denoising retained most of the reads for both loci (≥98%). For ITS, chimeras were identified in ∼2% of the sequence reads but <1% for ITS-28S sequence reads. At the end of the quality filtering procedure, 58% of the reads were retained for the ITS and 45% for the ITS-28S.). The final sequencing depth was 1190 reads or more for 75% of the samples for the ITS but only 112 reads or more for 75% of the samples for the ITS-28S. Hence, the ITS amplicon produced significantly more high-quality reads after the filtering steps. The two target amplicons differed in total sequencing yield and proportion of reads kept after the quality filtering steps (Figure 2C and 2D). Successfully identifying both primer sequences was a critical step for read retention during the ITS-28S filtering process. The ITS-28S amplicons showed also more variability among samples in retained read proportions compared to the ITS (Figure 3A-C). Samples with more reads showed a weak tendency to have a higher proportion of kept reads (R=0.57, Figure 3B). This tendency is stronger for ITS-28S amplicon (R=0.66). To better understand the shifts between the two datasets, we tested multiple filter parameter values for their impact on read retention. However, more relaxed primer matching parameters did not meaningfully increase retained reads. This is true for up to six allowed mismatches for primer detection. Beyond six allowed mismatches, a higher proportion of reads were kept as expected (Figure 3D). In summary, the difference in retained reads between amplicon datasets is largely unrelated to the primer matching stringency.

**Figure 3:**
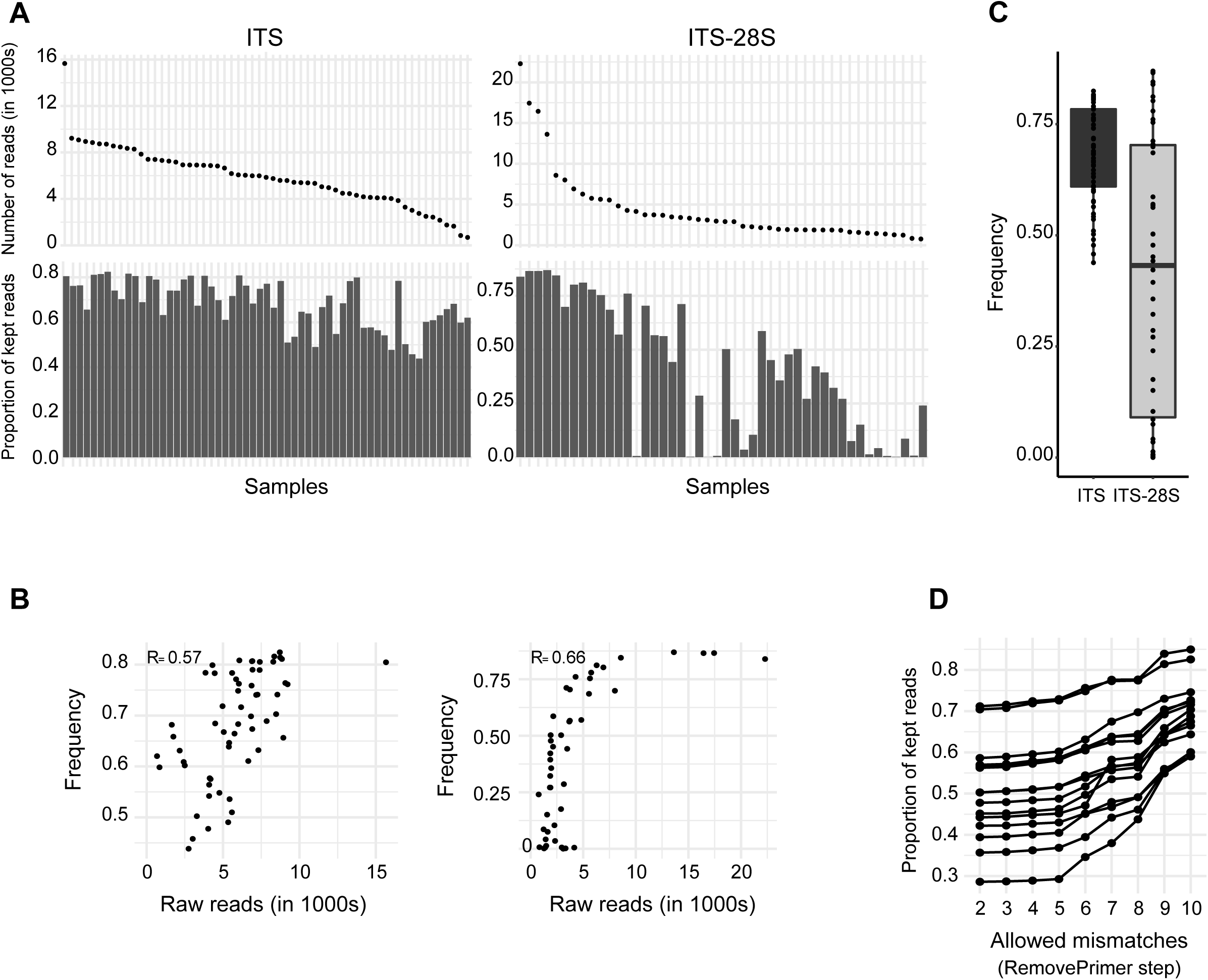
Impact of read filtering steps. A) Samples ranked by raw read counts and proportion of kept reads by sample at the primer trimming step for ITS and ITS-28S. B) Relation between raw read counts and proportion of kept reads at the primer trimming step for ITS and ITS-28S. C) Proportion of retained reads at the primer trimming step for both markers. D) Proportion of kept reads according to the number of allowed mismatches in primers detection for ITS-28S.

### Comparison of sequence variant recovery for the two barcoding loci

The number of inferred amplified sequence variants (ASVs) is highly variable among samples for both loci (Figure 4A). We found only a moderate correlation between the number of reads and the ASVs inferred per samples (R=0.39 for ITS and R=0.53 for ITS-28S, Figure 4B). For ITS we found a strong density peak of 500 reads by unique ASVs with a normal distribution. For the ITS-28S dataset, the distribution of reads by ASVs is more variable with a flatter curve going from 50 to 1000 reads by unique ASVs (Figure 4C). Overall, 933 ASVs were detected including both datasets. The ITS dataset comprises 888 ASVs among 59 samples and 175 ASVs among 46 samples for the ITS-28S dataset. The artificially cut ITS-28S fragment covers still 164 ASVs for the ITS subset and 140 for the 28S subset (Figure 4D). The sub-setting of the long fragment provides a direct comparison of the represented sequences in ITS and ITS-28S amplicon dataset, respectively. In a direct comparison, we found 888 ASVs with the ITS, 164 ASVs with ITS-28S amplicon subset to the ITS of which 119 (12.7%) were shared among the two amplicon datasets (Figure 5A). A total of 45 ASVs were detected only by the ITS-28S subset to the ITS amplicon and 769 ASVs were detected only by ITS amplicon. Among the shared ASVs, the proportion occupied by each regarding the two datasets are very similar. Overall, 83% of the shared ASVs differ by less than 1% in relative proportion between the two amplicon sets (Figure 5B). The most differentiated ASVs in terms of relative proportion are ASVs assigned to *Fomitiporia punctata* (8% more abundant in the ITS-28S subset to ITS and *Bacidina neosquamulosa* (4% more present in the ITS). Hence, the long fragment reveals only a minor degree of additional ASVs not already captured by the ITS amplicon.

**Figure 4:**
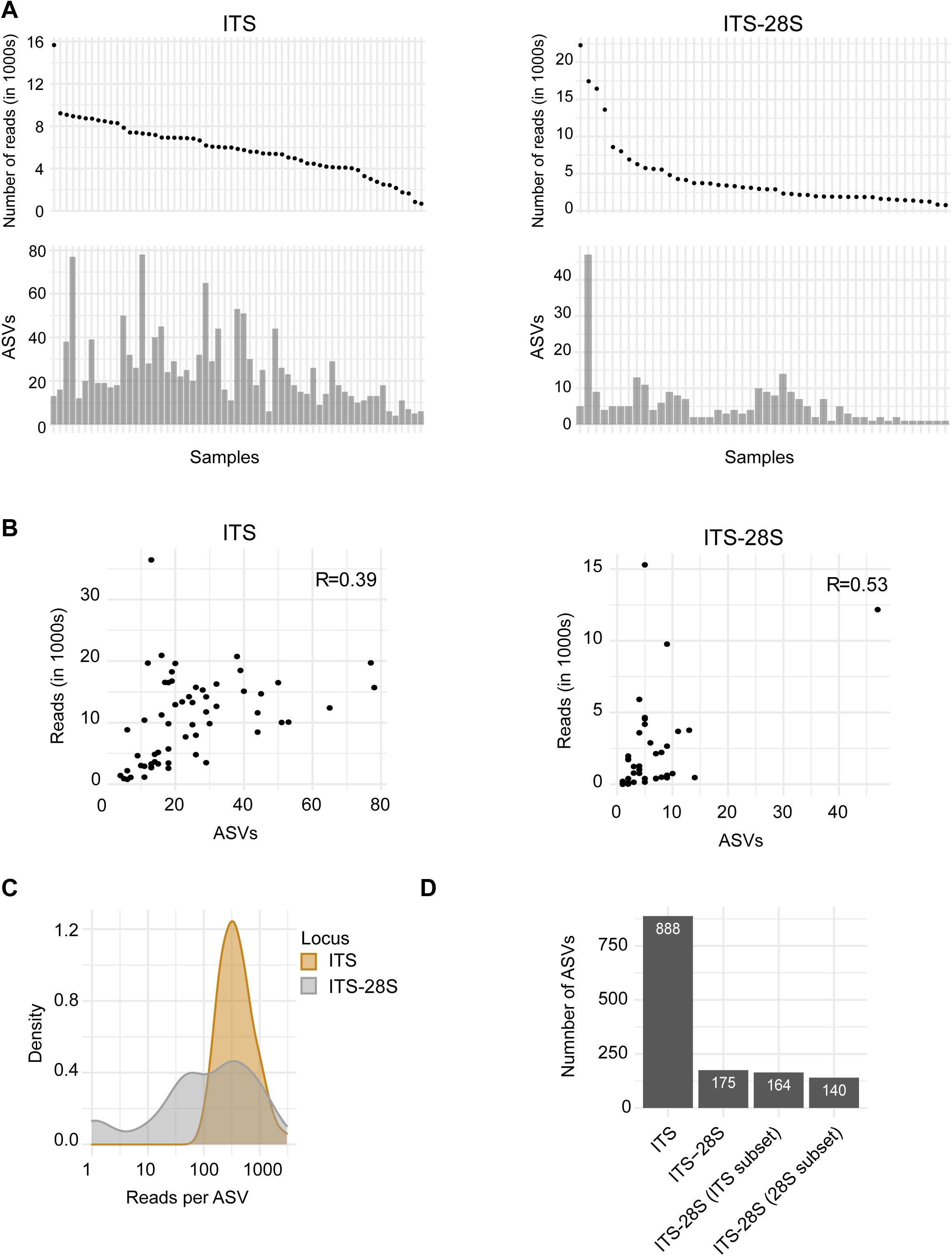
Amplified sequence variant (ASV) diversity. A) Samples ranked by raw read counts and inferred ASVs by sample for ITS and ITS-28S. B) Relation between raw read counts and ASVs detected for ITS and ITS-28S. C) Distribution of reads by ASVs for ITS and ITS-28S. D) ASVs detected by markers.

**Figure 5:**
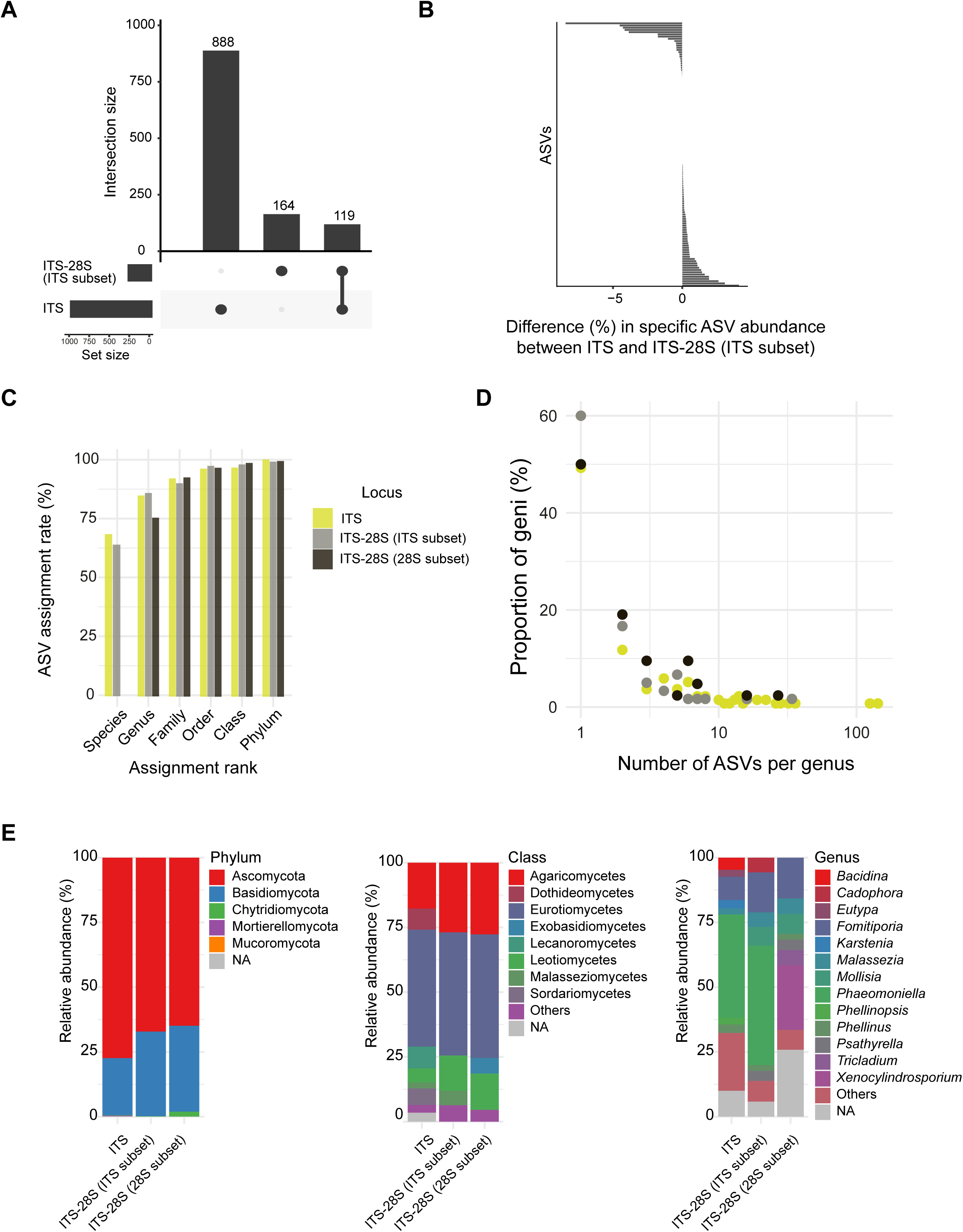
Taxonomical diversity among grapevine trunk samples. A) Intersecting sets of ASVs from ITS and ITS-28S ITS subset dataset. B) Proportional divergence of the shared ASVs of ITS and ITS-28S ITS subset. C) Proportion of assigned reads at several taxonomical ranks. D) Proportion of geni by number of ASVs for ITS, ITS-28S ITS subset and ITS-28S 28S subset. E) Relative abundance of several taxonomical ranks (phylum, class, genus) for ITS, ITS-28S ITS subset and ITS-28S 28S subset.

### Community composition of the grapevine mycobiome

To better understand the consequences of targeting either the ITS or the ITS-28S on the detected fungal community composition, we examined the diversity present in both datasets across several taxonomical ranks. In a first analysis, we subset the ITS-28S amplicon to more conserved LSU portion. This subset of the ITS-28S typically provided only genus-level resolution using the RDP database. For the ITS portion of the ITS-28S amplicon, as well as the ITS amplicon, ≥60% of ASVs were assigned at the species level (Figure 5C). Next, we analyzed the relative abundance of reads assigned to each phylum. Ascomycota represented the highest proportion of the reads (77%) in the ITS dataset and slightly less in the ITS-28S ITS subset (67%) and 28S subset (65%). Basidiomycota proportions showed opposite patterns with the ITS dataset showing 22%, ITS-28S ITS subset (33%) and 28S subset (33%; Figure 5E). For classes represented by more than 2% of the reads, we identified Eurotiomycetes and Agaricomycetes as the most abundant in all datasets. Dothideomycetes, Sordariomycetes and Lecanoromycetes are represented by >2% of the reads only in the ITS dataset. Similarly, Exobasidiomycetes were only found in the ITS-28S 28S subset at ≥2% (Figure 5E). The most represented genera according to the ITS amplicon were *Phaeomoniella* (40%), *Fomitiporia* (9%) and *Bacidina* (4.6%). For the ITS-28S ITS subset, the genera were similarly *Phaeomoniella* (46%), *Fomitiporia* (15%) and *Mollisia* (7%). For the ITS-28S 28S subset, we found *Xenocylindrosporium* (25%), *Fomitiporia* (16%) and *Mollisia* (8%) (Figure 5E).

At the species level, the diversity detected for the ITS-28S ITS subset was 45 ASVs. These ASVs correspond to 17 taxa with around half (*n* = 8) of the species or genera unable to be detected with ITS marker (Table supplementary material). The taxa detected only by the ITS subset of the ITS-28S amplicon were *Mollisia* sp. (8301 sequences), *Pseudoophiobolus rosae* (49 seq), *Meyerozyma guilliermondii* (19 seq), *Coniochaeta coluteae* (9 seq), *Bipolaris drechsleri* (6 seq), *Seimatosporium pistaciae* (6 seq), *Wojnowiciella cissampeli* (3 seq) and the Ceratobasidiaceae family (2 seq). For the 769 ASVs detected only based on the ITS amplicon, 103 species correspond to species uniquely detected by the ITS amplicon. The most abundant species in the dataset typically included multiple distinct ASVs matching to the same species. A total of 50% (ITS, ITS-28S 28S subset) and 60% (ITS-28S ITS subset) of the detected genera were represented by unique ASVs. Around 18% (ITS and ITS-28S ITS subset) and 16% (ITS-28S 28S subset) of the genera were represented by three distinct ASVs (Figure 5D).

The species represented by the highest number of ASVs is *P. chlamydospora* for both datasets. This species is represented by 304 (ITS) and 75 (ITS-28S ITS subsets) different ASVs highlighting significant intra-specific variation. *P. chlamydospora* is also the most abundant species in the ITS and ITS-28S ITS subset datasets. The species, which is typically associated with grapevine trunk disease was identified on 88% (ITS) and 97% (ITS-28S ITS subset) of the sampled plant. *P. chlamydospora* was not detected in the ITS-28S 28S subset. As this was surprising, we independently analyzed assignments of some ASVs using BLAST searches on NCBI GenBank. Some ASVs classified as belonging to the *Xenocylindrosporium* genus on the basis of matches in the RDP database rather belong to the *Phaeomoniella* genus based on GenBank matches. Uncertainty about taxonomic assignments using RDP is consistent with concerns about representativeness issues of long ribosomal fragment databases.

## Discussion

We used PacBio long-read technology to amplify and sequence two fungal barcoding amplicons (ITS and ITS-28S) in parallel from DNA extracted from a set of vine wood samples. Using identical handling, amplification and sequencing procedures, we obtained higher sequencing depth and higher ASV diversity for the shorter amplicon (*i*.*e*., ITS). We found no meaningful bias in the phylogenetic representation of the samples according to the selected amplicon. Despite the increased resolution of the long ITS-28S amplicon, the higher and more consistent yields of the shorter amplicons produced a clearer resolution of the fungal community of grapevine stems.

### Recovery of fungal barcoding sequences

The PacBio sequencing libraries prepared in parallel for the two different ribosomal amplicons differed in yield with ∼1.5x more sequences obtained for the ITS. Similarly, quality filtering retained a higher proportion of sequencing reads from the shorter amplicon. The higher yield and quality was consistent with findings by Tedersoo et al. (26). Differences in the number of sequences recovered for the two amplicons could be due to less efficient PCR amplification, *e*.*g*., due to competition amongst primed amplicons during the adaptor ligation step (26). And also because LR5 might be less efficient than ITS4 to amplify fungal diversity (33). Longer amplicons can also show reduced yields due to template positioning in sequencing wells (24, 26). Difficulties in properly detecting primer sequences on the long fragment could be due to a deterioration of sequencing quality. However, circular consensus sequencing should produce homogeneous quality scores for the entire template. Technological progress with the Sequel I system of PacBio and library preparation overall likely benefited read retention during quality filtering. Both our amplicons showed higher retention (58% for ITS and 45% for ITS-28S) compared to Tedersoo et al. (26) (28% of retained reads with a Sequel I system for a 400-700bp amplicon and 24% with a RSII system for a 1250-1700bp amplicon. With further progress, sequencing of longer fragments should become even more efficient.

### Contrasting the recovered sequence diversity

Targeting either a shorter amplicon with higher sequencing depth or a longer amplicon with reduced depth creates a trade-off that needs to be resolved. Both longer fragments and higher depth have the potential to improve the resolution of taxa present in a sample. We detected more ASVs using the ITS dataset, but we found no clear relationship between the sequencing depth and detected diversity as expected. As our study covered environmental samples, the sequence diversity is most likely highly heterogeneous. Hence, sequencing depth versus sequence diversity correlations among samples are only of limited use. Indeed, we detected samples of intermediate sequencing depth but with some of the highest numbers of recovered ASVs. Indeed, above some threshold, increasing sequencing depth is not expected to yield more recovered diversity (27). Purahong et al. (21) showed also that sequencing depth alone is a poor performance metric to evaluate the representation of the fungal community. Kennedy et al. (27) suggested a threshold of 100 reads per sample beyond which additional reads are unlikely to substantially shift community assessment of environmental samples. In our study, we obtained >1190 reads for the ITS and >112 reads for the ITS-28S amplicons for 75% of the samples. Hence, regardless of the selected amplicon, fungal communities should be reasonably well assessed in our study.

A basic argument for preferring longer amplicon sequences is the ability to detect a larger number of sequence variants present in the datasets (*i*.*e*., ASVs). Interestingly, our study recovered only a few ASVs using the longer ITS-28S that were not detected by the shorter amplicon (45 ASVs). It is likely that we have somewhat underestimated ASV diversity based on the shorter amplicon due to the chosen length cut-offs. Hence, using ITS is more beneficial to assess the fungal diversity present in the analysed grapevine trunk samples than targeting a longer amplicon making use of the highly accurate PacBio consensus reads. Some studies have even reported that increasing target amplicon lengths has negative effects on the assessment of microbial richness and community composition (34, 35). Our own comparative analyses have not revealed such detrimental effects. ASVs recovered both with ITS and ITS-28S amplicons showed very similar relative abundances among amplicons.

### Recovery of taxonomical diversity

We compared how well amplicons could be assigned to different taxonomical levels depending on the marker targeted. Taxonomic identification was similar for the ITS dataset compared to the ITS sequence extracted from the longer amplicon as well as for the entire ITS-28S amplicon (>75% of the ASVs assigned to the genus level for the three amplicon datasets). A somewhat lower proportion of the 28S sequences extracted from the ITS-28S amplicon were assigned, which is most likely explained by the poor taxon coverage of the 28S, compared to ITS, in sequence databases (∼15x more ITS sequences available in UNITE). The generally high levels of taxonomic assignments rates are consistent with the general growth of amplicon databases and the high accuracy of consensus sequences compared to previous studies relying on short read sequencing (*i*.*e*., Illumina). Furthermore, improvements in the analysis’s pipelines (*i*.*e*., DADA2) reduced erroneous chimera sequences and increased the accuracy of amplicon datasets. Our parallel diversity analyses based on two ribosomal amplicons revealed highly diverse fungal communities across grapevine trunks sampled across a vineyard. Our findings are consistent with previous analyses by Del Frari et al. (30) and Travadon et al. (36). We found high consistency in the identified taxa (8 out of 14 geni identified by Del Frari (30)). The consistently recovered taxa include *Phaeomoniella chlamydospora, Phaeoacremonium* spp., *Fomitiporia mediterranea*, fungal species commonly associated to esca disease of grapevine (37). This is consistent with expectations for fungal diversity sampled at the grafting point. Our implementation of PacBio amplicon sequencing opens up opportunities for high resolution profiling across large sets of samples covering space and time in mature vineyards. The high precision of the recovered sequences will allow the monitoring of fungal strains associated with GTD and more generally of wood endophyte fungal species.

A limitation in our comparison lies though in the lack of truly comparable reference databases for taxonomic assignments. Excising the ITS portion of the longer amplicon allowed for a direct comparison using the same database but a full comparison between the shorter and longer amplicon would require a fully equivalent ITS-28S sequence database. Our analyses match findings by Brown et al. (38) and Porras-Alfaro et al. (39) similarly comparing ITS and 28S amplicon diversity. It is evident that progress in exploiting longer amplicons to their full potential will require more comprehensive databases commensurate with the progress of sequencing technology.

## Material and methods

### Sample collection

Grapevine trunk sampling was conducted in La Côte vineyards in Echichens, Switzerland (46°32′07.034″N 6°30′03.807″E, WGS 84, 475m above sea). Samples were collected from 60 different plants in a single vineyard plot of 150m x 40m. Grapevine plants were all of the variety Gamaret (a cross between Gamay and Reichensteiner) grafted onto 3309C rootstock (*V. riparia x V. rupestris*). All the plants came from the same nursery (Dutruy in Founex, Switzerland) when there were planted in 2003. Plants affected by GTD had been replaced continuously and were not considered in our sampling. A plant was considered diseased if one or more shoots showed leaf necrosis up to wilt or dieback symptoms. These are typically symptoms classified as grapevine trunk disease (4, 40, 41). Grapevine wood was sampled at the grafting point (where fungal diversity was previously shown to be highest) using a non-destructive method. A 0.5 cm^2^ of bark was removed with a surface sterilized (EtOH 80%) scalpel. The sampling was then performed with a power drill with a surface sterilized drill bit (Ø 3.5 mm) by running the drill gently where the bark was removed to collect the coiled wood (∼60 mg) in an Eppendorf tube held underneath with sterilized tweezers. In the Eppendorf tubes, kept in an ice box during the sampling process, two 5 mm iron beads were previously deposited to ease the next step of the protocol. As soon as possible, the Eppendorf tubes containing the coiled wood were stored in -80°C (Figure 1).

### DNA extraction from wood samples

Eppendorf tubes (safe lock) containing two iron 5 mm beads and wood samples were taken out of the -80°C freezer and put in liquid nitrogen. The Eppendorf tubes were then placed two times for 1 min at 30 Hz in a TissueLyser (Qiagen Inc., Germantown, MD, USA) to disrupt wood tissues. Between and after these two steps of tissue disruption, tubes were placed in liquid nitrogen for 1 min. After having placed the tubes on ice to let them gently unfreeze, 1 mL CTAB was poured in each tube. The samples were then centrifuged for 1 min at 15’000 rounds/min and the supernatant was transferred to a new tube. The fungal DNA extraction was then performed with phenol-chloroform as described in Hofstetter et al. (42). The DNA quality was checked with an electrophoresis gel and the extracted products were stored at - 80°C.

### Amplification of fungal ribosomal DNA

Two loci were targeted for amplification: ITS using primers ITS1F and ITS4 and a longer amplicon including ITS and a portion of the 28S using primers ITS1F and LR5 (Figure 1). We followed the PacBio procedure using barcoded universal primers for multiplexing amplicons, which includes two PCR steps (see https://www.pacb.com). The first PCR program was 30 sec of denaturation at 98°C, then 30 cycles of 15 sec at 98°C, 15 sec at 55°C and 1:30 min at 72°C, followed by a final elongation step for 7 min at 72°C. The second PCR program was 30 sec of denaturation at 98°C, then 20 cycles of 15 sec at 98°C, 15 sec at 64°C and 1:20 min at 72°C, followed by a final elongation step for 7 min at 72°C. We performed purification between the two PCRs to reduce contaminants or carry-over of primer dimers using 96-well PCR purification plates (Qiagen Inc., Germantown, MD, USA). The final libraries were quantified with a Qubit Fluorometer (ThermoFischer, Foster City, CA, USA), then all samples were pooled equimolarly. The pooled samples were then purified with 1X AMPure XP Beads (Beckman Coulter Inc., Indianapolis, USA) as per the manufacturer’s instructions. Amplicons were prepared for SMRT sequencing at the Functional Genomics Center in Zürich (FGCZ), Switzerland. Sequencing was performed on the PacBio Sequel II platform.

### Demultiplexing and trimming

The raw reads were demultiplexed using lima. After obtaining fastq files, reads were processed for quality filtering with the DADA2 package for R (Figure 1, https://github.com/benjjneb/dada2). The DADA2 processing steps include: i) dereplication with primer detection. Reads without primer sequences are discarded; ii) filtering for amplicon length (between 500-1000 bp for ITS and 1000-2000 for the longer fragment); iii) error detection where error rates were learnt by alternating between sample inference and error rate estimation until convergence. A feature table of observed transitions for each type and quality scores were produced; iv) denoising to reduce sequencing errors based on error models; v) checking for chimeras where each sequence is evaluated against a set of putative parental sequences drawn from the sequence collection. Several tests were performed with removeBimeraDenovo by increasing the minFoldParentOverAbundance parameter from 4 to 8. We chose to continue with minFoldParentOverAbundance = 8 to retain a maximum of reads.

### Analyses of amplicon sequence variants and taxonomic assignments

We used the DADA2 algorithm to infer amplicon sequence variants (ASVs) from the filtered reads (Figure 1). ASVs represent reads with 100% similarity accounting for sequencing errors by appropriately modelling PacBio CCS sequencing errors (18). Taxonomic assignments were performed with the function AssignTaxonomy of DADA2 pipeline which classifies sequences based on reference training datasets. The databases used for assignments were UNITE (https://unite.ut.ee/cite.php) for the ITS and the Ribosomal Database Project (RDP) (RDP LSU training set). To compare sequence diversity directly between the short and long amplicon sets, ITS-28S fragments were cut in two subset fragments corresponding to the ITS and the28S (beginning of LSU), respectively. This was performed using the removePrimers step in the DADA2 pipeline. Instead of detecting ITS1F-LR5 primer pairs as above, we used ITS1F-ITS4 for the ITS and rcITS4-LR5 for the 28S. The sequence subsetting produced the two subsets ITS-28S subset to ITS and ITS-28S subset to 28S. All further analyses were performed with R (v4.0.3) (R Core Team, 2020).

## Acknowledgements

Library preparation and sequencing was performed at the Functional Genomics Centre Zurich (FGCZ). We are grateful to the members of the mycology research group of Agroscope (N. Lecoultre, E. Michellod, A.-L. Fabre, P.-H. Dubuis, D. Restori, A. Melgar) for their assistance in sampling vineyards.

## References

1. Nilsson RH, Anslan S, Bahram M, Wurzbacher C, Baldrian P, Tedersoo L. 2019. Mycobiome diversity: high-throughput sequencing and identification of fungi. Nat Rev Microbiol 17:95–109.

2. Fiorilli V, Catoni M, Lanfranco L, Zabet NR. 2020. Editorial: Interactions of Plants With Bacteria and Fungi: Molecular and Epigenetic Plasticity of the Host. Front Plant Sci 11:274.

3. Yadeta KA, Thomma BPHJ. 2013. The xylem as battleground for plant hosts and vascular wilt pathogens. Front Plant Sci 4:97.

4. Bertsch C, Ramírez-Suero M, Magnin-Robert M, Larignon P, Chong J, Abou-Mansour E, Spagnolo A, Clément C, Fontaine F. 2013. Grapevine trunk diseases: complex and still poorly understood. Plant Pathol 62:243–265.

5. Agrios, G. (2005) Plant Pathology. 5th Edition, Elsevier. https://www.scirp.org › reference › r..https://www.scirp.org › reference › r.

6. Pouzoulet J, Pivovaroff AL, Santiago LS, Rolshausen PE. 2014. Can vessel dimension explain tolerance toward fungal vascular wilt diseases in woody plants? Lessons from Dutch elm disease and esca disease in grapevine. Front Plant Sci 5:253.

7. Vieites JM, Guazzaroni M-E, Beloqui A, Golyshin PN, Ferrer M. 2009. Metagenomics approaches in systems microbiology. FEMS Microbiol Rev 33:236–255.

8. Tiew PY, Mac Aogain M, Ali NABM, Thng KX, Goh K, Lau KJX, Chotirmall SH. 2020. The Mycobiome in Health and Disease: Emerging Concepts, Methodologies and Challenges. Mycopathologia 185:207–231.

9. Zhou J, He Z, Yang Y, Deng Y, Tringe SG, Alvarez-Cohen L. 2015. High-throughput metagenomic technologies for complex microbial community analysis: open and closed formats. MBio 6.

10. O’Brien HE, Parrent JL, Jackson JA, Moncalvo J-M, Vilgalys R. 2005. Fungal community analysis by large-scale sequencing of environmental samples. Appl Environ Microbiol 71:5544–5550.

11. Wu B, Hussain M, Zhang W, Stadler M, Liu X, Xiang M. 2019. Current insights into fungal species diversity and perspective on naming the environmental DNA sequences of fungi. Mycology 10:127–140.

12. Shokralla S, Porter TM, Gibson JF, Dobosz R, Janzen DH, Hallwachs W, Golding GB, Hajibabaei M. 2015. Massively parallel multiplex DNA sequencing for specimen identification using an Illumina MiSeq platform. Sci Rep 5:1–7.

13. Heeger F, Wurzbacher C, Bourne EC, Mazzoni CJ, Monaghan MT. 2019. Combining the 5.8S and ITS2 to improve classification of fungi. Methods Ecol Evol 10:1702–1711.

14. Schoch CL, Seifert KA, Huhndorf S, Robert V, Spouge JL, Levesque CA, Chen W, Null N, Null N, Bolchacova E, Voigt K, Crous PW, Miller AN, Wingfield MJ, Aime MC, An K-D, Bai F-Y, Barreto RW, Begerow D, Bergeron M-J, Blackwell M, Boekhout T, Bogale M, Boonyuen N, Burgaz AR, Buyck B, Cai L, Cai Q, Cardinali G, Chaverri P, Coppins BJ, Crespo A, Cubas P, Cummings C, Damm U, de Beer ZW, de Hoog GS, Del-Prado R, Dentinger B, Diéguez-Uribeondo J, Divakar PK, Douglas B, Dueñas M, Duong TA, Eberhardt U, Edwards JE, Elshahed MS, Fliegerova K, Furtado M, García MA, Ge Z-W, Griffith GW, Griffiths K, Groenewald JZ, Groenewald M, Grube M, Gryzenhout M, Guo L-D, Hagen F, Hambleton S, Hamelin RC, Hansen K, Harrold P, Heller G, Herrera C, Hirayama K, Hirooka Y, Ho H-M, Hoffmann K, Hofstetter V, Högnabba F, Hollingsworth PM, Hong S-B, Hosaka K, Houbraken J, Hughes K, Huhtinen S, Hyde KD, James T, Johnson EM, Johnson JE, Johnston PR, Jones EBG, Kelly LJ, Kirk PM, Knapp DG, Kõljalg U, Kovács GM, Kurtzman CP, Landvik S, Leavitt SD, Liggenstoffer AS, Liimatainen K, Lombard L, Luangsa-ard JJ, Lumbsch HT, Maganti H, Maharachchikumbura SSN, Martin MP, May TW, McTaggart AR, Methven AS, Meyer W, Moncalvo J-M, Mongkolsamrit S, Nagy LG, Nilsson RH, Niskanen T, Nyilasi I, Okada G, Okane I, Olariaga I, Otte J, Papp T, Park D, Petkovits T, Pino-Bodas R, Quaedvlieg W, Raja HA, Redecker D, Rintoul TL, Ruibal C, Sarmiento-Ramírez JM, Schmitt I, Schüßler A, Shearer C, Sotome K, Stefani FOP, Stenroos S, Stielow B, Stockinger H, Suetrong S, Suh S-O, Sung G-H, Suzuki M, Tanaka K, Tedersoo L, Telleria MT, Tretter E, Untereiner WA, Urbina H, Vágvölgyi C, Vialle A, Vu TD, Walther G, Wang Q-M, Wang Y, Weir BS, Weiß M, White MM, Xu J, Yahr R, Yang ZL, Yurkov A, Zamora J-C, Zhang N, Zhuang W-Y, Schindel D. 2012. Nuclear ribosomal internal transcribed spacer (ITS) region as a universal DNA barcode marker for Fungi. Proceedings of the National Academy of Sciences 109:6241–6246.

15. Bálint M, Schmidt P-A, Sharma R, Thines M, Schmitt I. 2014. An Illumina metabarcoding pipeline for fungi. Ecol Evol 4:2642–2653.

16. Kõljalg U, Nilsson RH, Abarenkov K, Tedersoo L, Taylor AFS, Bahram M, Bates ST, Bruns TD, Bengtsson-Palme J, Callaghan TM, Douglas B, Drenkhan T, Eberhardt U, Dueñas M, Grebenc T, Griffith GW, Hartmann M, Kirk PM, Kohout P, Larsson E, Lindahl BD, Lücking R, Martín MP, Matheny PB, Nguyen NH, Niskanen T, Oja J, Peay KG, Peintner U, Peterson M, Põldmaa K, Saag L, Saar I, Schüßler A, Scott JA, Senés C, Smith ME, Suija A, Taylor DL, Telleria MT, Weiss M, Larsson K-H. 2013. Towards a unified paradigm for sequence-based identification of fungi. Mol Ecol 22:5271–5277.

17. Schlaeppi K, Bender SF, Mascher F, Russo G, Patrignani A, Camenzind T, Hempel S, Rillig MC, van der Heijden MGA. 2016. High-resolution community profiling of arbuscular mycorrhizal fungi. New Phytol 212:780–791.

18. Callahan BJ, Wong J, Heiner C, Oh S, Theriot CM, Gulati AS, McGill SK, Dougherty MK. 2019. High-throughput amplicon sequencing of the full-length 16S rRNA gene with single-nucleotide resolution. Nucleic Acids Res 47:e103.

19. Hebert PDN, Braukmann TWA, Prosser SWJ, Ratnasingham S, deWaard JR, Ivanova NV, Janzen DH, Hallwachs W, Naik S, Sones JE, Zakharov EV. 2018. A Sequel to Sanger: amplicon sequencing that scales. BMC Genomics 19:219.

20. Tedersoo L, Anslan S. 2019. Towards PacBio-based pan-eukaryote metabarcoding using full-length ITS sequences. Environ Microbiol Rep 11:659–668.

21. Purahong W, Mapook A, Wu Y-T, Chen C-T. 2019. Characterization of the Castanopsis carlesii Deadwood Mycobiome by Pacbio Sequencing of the Full-Length Fungal Nuclear Ribosomal Internal Transcribed Spacer (ITS). Front Microbiol 10:983.

22. Jiao X, Zheng X, Ma L, Kutty G, Gogineni E, Sun Q, Sherman BT, Hu X, Jones K, Raley C, Tran B, Munroe DJ, Stephens R, Liang D, Imamichi T, Kovacs JA, Lempicki RA, Huang DW. 2013. A Benchmark Study on Error Assessment and Quality Control of CCS Reads Derived from the PacBio RS. J Data Mining Genomics Proteomics 4.

23. Larsen PA, Heilman AM, Yoder AD. 2014. The utility of PacBio circular consensus sequencing for characterizing complex gene families in non-model organisms. BMC Genomics 15:720.

24. Castaño C, Berlin A, Brandström Durling M, Ihrmark K, Lindahl BD, Stenlid J, Clemmensen KE, Olson Å. 2020. Optimized metabarcoding with Pacific biosciences enables semi-quantitative analysis of fungal communities. New Phytol 228.

25. Singer E, Bushnell B, Coleman-Derr D, Bowman B, Bowers RM, Levy A, Gies EA, Cheng J-F, Copeland A, Klenk H-P, Hallam SJ, Hugenholtz P, Tringe SG, Woyke T. 2016. High-resolution phylogenetic microbial community profiling. ISME J 10:2020–2032.

26. Tedersoo L, Tooming-Klunderud A, Anslan S. 2018. PacBio metabarcoding of Fungi and other eukaryotes: errors, biases and perspectives. New Phytol 217:1370–1385.

27. Kennedy PG, Cline LC, Song Z. 2018. Probing promise versus performance in longer read fungal metabarcoding. New Phytol.

28. Fuks G, Elgart M, Amir A, Zeisel A, Turnbaugh PJ, Soen Y, Shental N. 2018. Combining 16S rRNA gene variable regions enables high-resolution microbial community profiling. Microbiome 6:17.

29. Edgar RC. 2018. Updating the 97% identity threshold for 16S ribosomal RNA OTUs. Bioinformatics 34:2371–2375.

30. Del Frari G, Gobbi A, Aggerbeck MR, Oliveira H, Hansen LH, Ferreira RB. 2019. Characterization of the Wood Mycobiome of Vitis vinifera in a Vineyard Affected by Esca. Spatial Distribution of Fungal Communities and Their Putative Relation With Leaf Symptoms. Front Plant Sci 10:910.

31. Hofstetter V, Buyck B, Croll D, Viret O, Couloux A, Gindro K. 2012. What if esca disease of grapevine were not a fungal disease? Fungal Divers 54:51–67.

32. Bruez E, Baumgartner K, Bastien S, Travadon R, Guérin-Dubrana L, Rey P. 2016. Various fungal communities colonise the functional wood tissues of old grapevines externally free from grapevine trunk disease symptoms. Aust J Grape Wine Res 22:288–295.

33. Asemaninejad A, Weerasuriya N, Gloor GB, Lindo Z, Thorn RG. 2016. New Primers for Discovering Fungal Diversity Using Nuclear Large Ribosomal DNA. PLoS One 11:e0159043.

34. Huber JA, Morrison HG, Huse SM, Neal PR, Sogin ML, Mark Welch DB. 2009. Effect of PCR amplicon size on assessments of clone library microbial diversity and community structure. Environ Microbiol 11:1292–1302.

35. Engelbrektson A, Kunin V, Wrighton KC, Zvenigorodsky N, Chen F, Ochman H, Hugenholtz P. 2010. Experimental factors affecting PCR-based estimates of microbial species richness and evenness. ISME J 4:642–647.

36. Travadon R, Rolshausen PE, Gubler WD, Cadle-Davidson L, Baumgartner K. 2013. Susceptibility of Cultivated and Wild Vitis spp. to Wood Infection by Fungal Trunk Pathogens. Plant Dis 97:1529–1536.

37. Surico G. 2009. Towards a redefinition of the diseases within the esca complex of grapevine. Phytopathol Mediterr 48:5–10.

38. Brown SP, Rigdon-Huss AR, Jumpponen A. 2014. Analyses of ITS and LSU gene regions provide congruent results on fungal community responses. Fungal Ecol 9:65–68.

39. Porras-Alfaro A, Liu K-L, Kuske CR, Xie G. 2014. From genus to phylum: large-subunit and internal transcribed spacer rRNA operon regions show similar classification accuracies influenced by database composition. Appl Environ Microbiol 80:829–840.

40. Mugnai L, Graniti A, Surico G. 1999. Esca (Black Measles) and Brown Wood-Streaking: Two Old and Elusive Diseases of Grapevines. Plant Dis 83:404–418.

41. Larignon P, Fontaine F, Farine S, Clément C, Bertsch C. 2009. Esca et Black Dead Arm : deux acteurs majeurs des maladies du bois chez la Vigne. C R Biol 332:765–783.

42. Hofstetter V, Clémençon H, Vilgalys R, Moncalvo J-M. 2002. Phylogenetic analyses of the Lyophylleae (Agaricales, Basidiomycota) based on nuclear and mitochondrial rDNA sequences. Mycol Res 106:1043–1059.

